# Spontaneous visual preference for face-like stimuli is impaired in newly-hatched domestic chicks exposed to valproic acid during embryogenesis

**DOI:** 10.1101/2021.04.12.436924

**Authors:** Alice Adiletta, Samantha Pedrana, Orsola Rosa-Salva, Paola Sgadò

## Abstract

One of the most fascinating properties of the human brain is the capacity of newborn babies to recognize and orient to faces and simple schematic face-like patterns since the first hours of life. A striking feature of these social orienting mechanisms is their transversal appearance in remarkably diverse vertebrate species. Similar to newborn babies, also non-human primates and domestic chicks have been shown to express orienting responses to faces and schematic face-like patterns. More importantly, existing studies have hypothesized that early disturbances of these mechanisms represent one of the earliest biomarkers of social deficits in autism spectrum disorders (ASD). Recent data suggest that newborns at high risk for the disorder express altered responses to schematic face-like configurations. Here we modeled ASD in domestic chicks using the anticonvulsant valproic acid (VPA), and tested the animals for their predisposed preference towards schematic face-like configuration stimuli. We found that VPA impairs the chicks’ preference responses to the social stimuli. Based on the results shown here and in previous studies, we propose the domestic chicks as elective animal models to study early-emerging neurobehavioural markers and to investigate the biological mechanisms underlying face processing deficits in ASD.

## Introduction

Biological predispositions to orient to and preferentially learn about conspecifics are one of the earliest expressions of social behavior in vertebrates and are critical for survival. These elementary behavioral markers of social orienting are spontaneous, possibly hard-wired, mechanisms that bias visual attention to simple features of animate beings since the earliest minutes of life (1, 2). Human faces and schematic face-like patterns generate remarkable responses in typical developing neonates (3). More strikingly, the same abilities can be observed in newly-hatched chicks (4, 5) and visually naïve monkeys (6, 7). Other species have also been shown to respond to similar schematic configurations (8), such that privileged face processing could be pervasive in vertebrates.

More importantly, it has been hypothesized that early disturbances of these social orienting mechanisms may be one of the earliest signs of social deficits in autism spectrum disorders (ASD), and might also contribute to the pathophysiology of these disorders by compromising, early on, the typical developmental trajectories of the social brain (9–12). In line with that, impairments in face and eye-gaze direction processing have been reported in infants at risk of ASD (13, 14) (but see Bradshaw et al., 2020 (15), Shultz et al., 2018 (16) and Jones and Klin, 2013 (17) for a critical discussion on impairments before 4-6 months of age).

Given the complexity of human social behaviour and the limitations that human studies impose, animal models are instrumental in providing clues on the nature and origin of these crucial social orienting mechanisms and their role in atypical social development. Valproic acid (VPA) exposure has been extensively used in several animal models to reproduce ASD core symptoms (18). Previous studies have shown that exposure to different doses of VPA during embryogenesis induces alterations of several aspects of social behaviour in domestic chicks (19, 20). We used VPA exposure to induce neurodevelopmental changes associated with social deficits in domestic chicks and tested whether VPA could impact the expression of early approach responses to schematic face-like patterns. We found that VPA impairs the chicks’ preference responses to these social stimuli. Based on the results shown here, we propose the domestic chicks as elective animal models to study these early-emerging neurobehavioural markers and to investigate the biological mechanisms underlying face processing deficits in ASD.

## Materials and Methods

### Ethical approval

All experiments were conducted according to the current Italian and European Community laws for the ethical treatment of animals. The experimental procedures were approved by the Ethical Committee of the University of Trento and licensed by the Italian Health Ministry (permit number 986/2016-PR).

### Embryo injections

Fertilized eggs of domestic chicks (*Gallus gallus*), of the Ross 308 (Aviagen) strain, were obtained from a local commercial hatchery (Agricola Berica, Montegalda (VI), Italy). Upon arrival the eggs were placed in the dark and incubated at 37.5 °C and 60% relative humidity, with rocking. One week before the predicted date of hatching, on embryonic day 14 (E14), fertilized eggs were selected by a light test, before injection. Chick embryo injection was performed according to previous reports (19, 21). Briefly, a small hole was made on the egg shell above the air sac, and 35 μmoles of VPA (Sodium Valproate, Sigma Aldrich) were administered to each fertilized egg, in a volume of 200 μl, by dropping the solution onto the chorioallantoic membrane (VPA group). Age-matched control eggs were injected using the same procedure with 200 μL of vehicle (double distilled injectable water; CTRL group). After sealing the hole with paper tape, eggs were placed back in the incubator until E18, when they were placed in a hatching incubator (FIEM srl, Italy). Hatching took place at a temperature of 37.7 °C, with 72% humidity. The day of hatching was considered post-hatching day 0 (P0).

### Rearing conditions

After hatching in darkness, 69 chicks (38 males and 31 females) were kept in the hatching incubator for 24 hours before the experiment.

### Apparatus and test stimuli

The test apparatus was a corridor, 45 cm long x 22.3 cm wide, made from wood and covered with opaque white plastic coating. The apparatus was divided in three sections (outlined on the apparatus floor), one central for positioning the animal, equidistant from the two stimuli, and two on the opposite side of the corridor, in proximity to the stimuli, considered the choice sectors. The stimuli were placed at the opposite side of the rectangular arena, on panels of light-filtering Plexiglas, lit by a 201 lumen LED placed behind the Plexiglas partition. The visual stimuli were previously described in Rosa-Salva et al. (2010) (4). Briefly, they consisted of featureless face silhouette shapes, made of orange stiff paper (10 x 5.6 cm, see Figure 1) that contained internal features: three black squares (of side 1 cm), organized as an upside down triangle for the schematic face-like configuration, or aligned vertically for the control nonsocial stimulus. Both stimuli were top-heavy configurations, having two elements in their upper part and one in their lower part.

**Figure 1.**
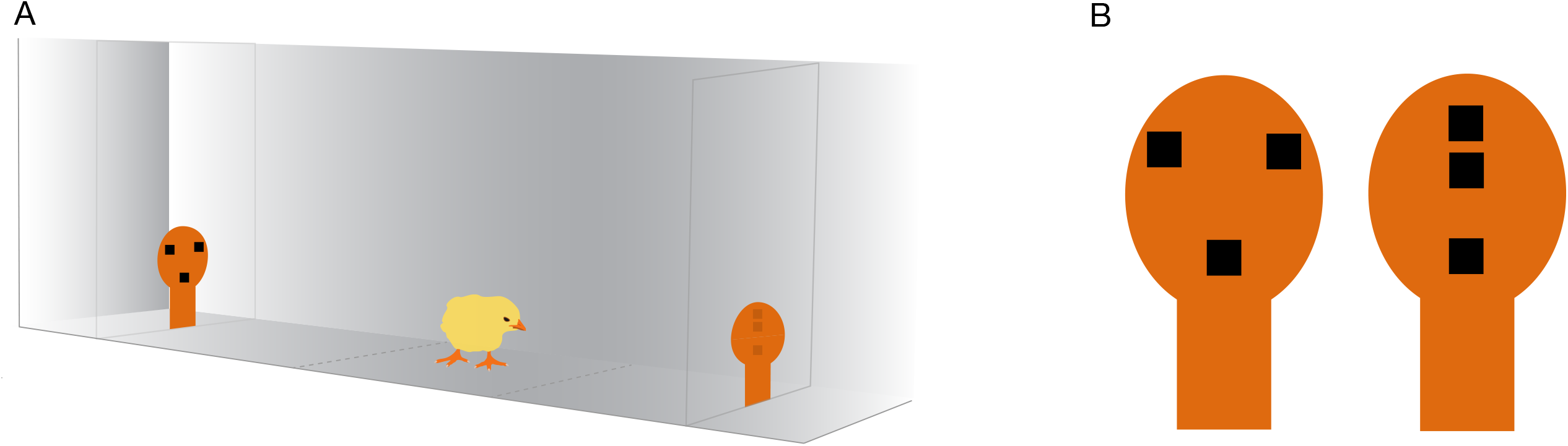
Schematic illustration of the social preference test apparatus and the stimuli. A, The chick was placed in the center of the arena and was free to approach either of the stimuli, placed at the two ends of the apparatus and lit by a 201 lumen LED. The chick’s behavior was video-recorded from above. B, The stimuli consisted of orange stiff paper silhouettes containing internal features resembling a face-like configuration (left) or a nonsocial control configuration (right). The chick image is courtesy of Openclipart (openclipart.org) under Creative Commons Zero 1.0 Public Domain License.

### Test procedures

At postnatal day 1 (P1), about 24 hours after hatching, chicks were transported in complete darkness to the test room and placed in the apparatus: positioning with respect to the test stimuli, as well as the left-right position of the stimuli in the apparatus, was counterbalanced across animals. The animals’ approach responses were recorded using a camera placed on top of the apparatus, for the entire duration of the test (12 minutes).

### Statistical analysis

The preference response was measured as a social preference index adjusted for the overall activity of the chicks during the test, calculated as the time spent in the choice sector close to the social stimulus (schematic face-like configuration) divided by the total time spent in the two choice sections (face-like + nonsocial). Values of this ratio range from 1 (full choice for the social stimulus) to 0 (full choice for the nonsocial stimulus), where 0.5 represents the absence of preference. Significant departures of the social preference index from chance level (0.5) were estimated by one-sample two-tailed t-tests. The number of chicks that first approached the two stimuli in the two treatment and gender groups was compared using one-sided Pearson’s chi-square test. We assessed differences in behavioural activity measuring the time required to move to one of the choice sections (latency to choice) and the number of sector switches (spontaneous alternations). Effect of Treatment and Sex on the social preference index, the latency to first choice and the spontaneous alternations was evaluated by multifactorial analysis of variance (ANOVA). Statistical analyses were performed with GraphPad Prism 9 and RStudio (package *CHAID* v 0.1-2). Alpha was set to 0.05 for all tests.

## Results

To assess the effect of VPA on face perception, and avoid any possible influence of previous experiences in evaluating the chicks’ approach to the stimuli, we excluded visual experience prior to the test. To obtain a better approach rate, we extended the duration of the test compared to the previous reports to 12 minutes. Using this adapted paradigm, we tested 69 chicks (31 females, 38 males), 24 hours after hatching. We found a significant difference between the treatment groups in the preference index for the schematic face-like configuration stimulus (Fig 2A; treatment: F_(1, 65)_ = 4.805, p = 0.0320; sex: F_(1, 65)_ = 0.5745, p = 0.4512; treatment*sex: F_(1, 65)_ = 2.652, p = 0.1083). While vehicle-injected chicks significantly prefered the schematic face-like stimulus, VPA-exposed chicks did not display any significant preference for this stimulus compared to what expected by chance (Fig 2A; CTRL t_(32)_ = 2.481, p = 0.0186; VPA t_(35)_ = 0.3425, p = 0.7341; group mean: CTRL 0.6694 [95% C.I. 0.5303-0.8085]; VPA 0.4764 [95% C.I. 0.3364-0.6164]). We then analyzed the latency to choice and the number of sector alternations after the first choice. We found a significant effect of treatment on the latency: VPA-injected chicks had a shorter latency to choice compared to controls (Fig 2B; treatment: F_(1, 65)_ = 5.369, p = 0.0237 sex: F_(1, 65)_ = 0.1881, p = 0.6660; treatment*sex: F_(1, 65)_ = 0.1270, p = 0.7228; group mean: CTRL 339 seconds [95% C.I. 275-403]; VPA 234 seconds [95% C.I. 172-295]). Spontaneous alternations in the two choice sectors did not significantly differ between treatment groups (Fig 2C; treatment: F_(1, 65)_ = 1.941, p = 0.1683; sex: F_(1, 65)_ = 0.0790, p = 0.7795; treatment*sex: F_(1, 65)_ = 1.293, p = 0.2598; group mean: CTRL 5.091 [95% C.I. 2.344-7.838]; VPA 8.389 [95% C.I. 5.077-11.70]).

**Figure 2.**
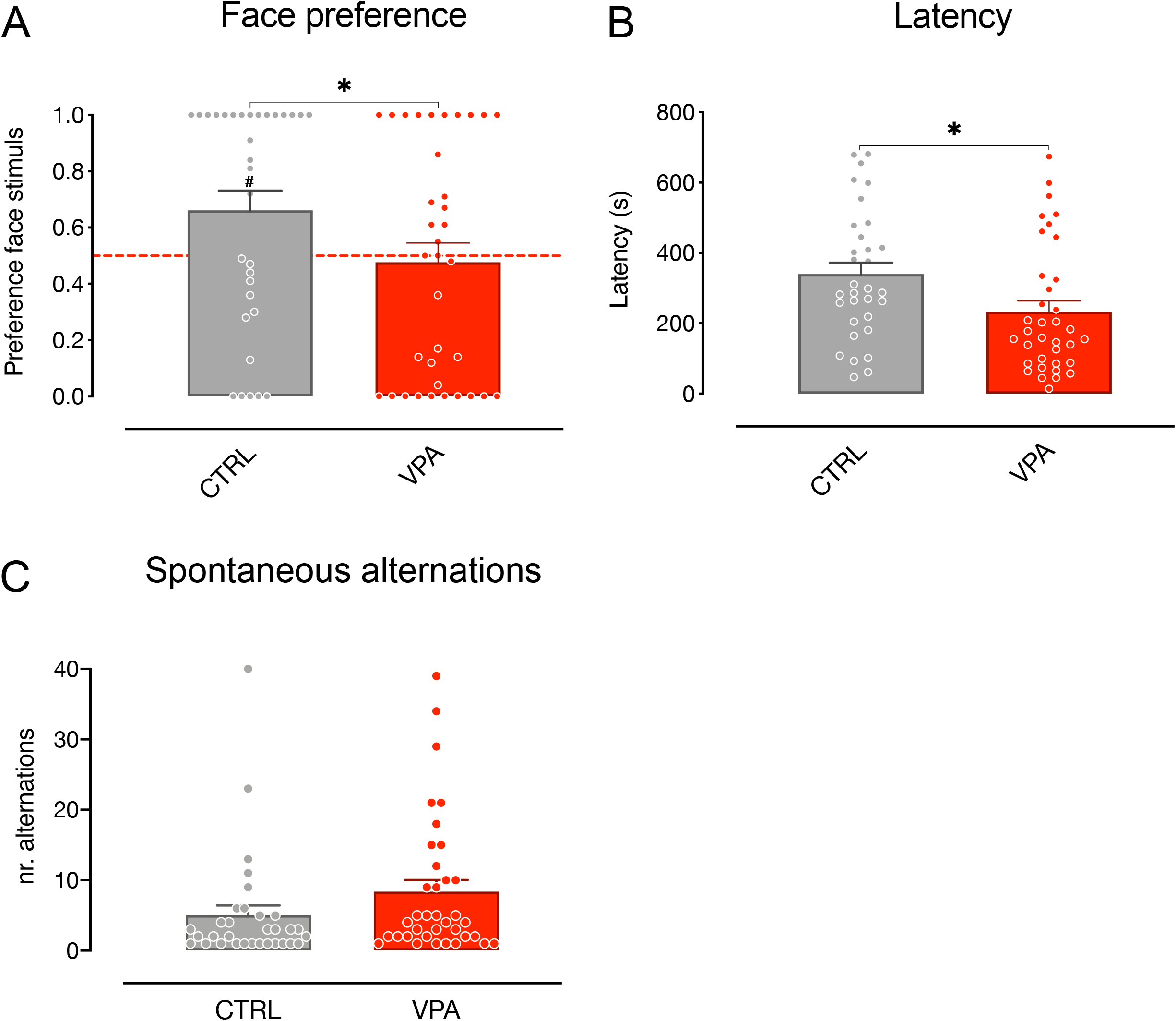
Spontaneous visual preference test. Bar graphs represent social preferences indexes (A), latency to first choice (B) and spontaneous alternations (C). A, social preference test for schematic face-like (social) stimulus and nonsocial stimulus (see Methods for details). Analysis of variance of social preference indexes using treatment and sex as between-subject factors, revealed a significant main effect of treatment and no other main effects or interactions among the factors analyzed. One-sample t-test on preference indexes indicate a significant difference from chance level for the control group, but not for VPA-treated chicks. The number sign (#) indicate significant departures of the preference index from chance level (0.5), marked by the red line. B, behavioural activity during the test measured as latency to choice. Analysis of variance on time taken by the chicks to move in one of the choice sections using treatment and sex as between-subject factors, showing a significant effect of treatment and no other main effects or interaction. C, behavioural activity during the test measured as sector switches. Analysis of variance on number of switches between the three sectors, using treatment and sex as between-subject factors, showing (C) no significant main effect of treatment or sex, and no interactions. Data represent Mean ± SEM, ^#^p < 0.05, *p < 0.05; **p < 0.01, ***p < 0.001.

The number of chicks that approached the face-like configuration as the first stimulus was not significantly different between treatment groups (one-sided Pearson’s ⍰ _1_ ^2^ = 2.944, p = 0.0862; CTRL: face N = 21, nonface N = 12, VPA: face N = 15, nonface N = 20, data not shown).

## Discussion

Newborns of several vertebrate species exhibit rudimental knowledge about the typical appearance of animate beings that orient the young organisms’ attention towards plausible social partners and caregivers. Several studies hypothesized that this mechanism contributes to create an early social bond with caretakers and social companions (9, 22), an essential process for subsequent social and language development. Newborn babies, as well as non-human primates and domestic chicks, have been shown to express remarkable orienting responses to faces and schematic face-like patterns (4–7). Divergence from these early social interactions may induce a cascade of maladaptive trajectories culminating in atypical social abilities, such as those observed in ASD.

Using the preference response to face-like stimuli as an evolutionarily conserved neurobehavioural marker and exploiting the advantages of animal models, we investigated whether these early-emerging social orienting mechanisms could be affected by a compound, VPA, known to interfere with development of the social brain. We examined the preference response towards schematic-face like configurations of animals whose pattern of brain development may have been altered by VPA, an anticonvulsant increasing the risk to develop ASD in humans. We found that VPA had a dramatic effect on the preference towards schematic-face configuration stimuli.

Previous studies have revealed a predisposed response to schematic face-like configurations in newly-hatched chicks, using both subjects imprinted on face-neutral stimuli and visually naïve subjects (4, 5, 23). To assess the effect of VPA on face perception, and avoid any possible influence of previous experiences in evaluating the chicks’ approach to the stimuli, we applied this latter experimental procedure, excluding visual experience prior to the test. Since dark reared animals are less active compared to chicks exposed to visual stimuli, to obtain a better approach rate, we extended the duration of the test compared to the previous reports. Increasing the test duration in our experiment contributed to heighten the approach response and the face preference, without introducing the potential influence of visual experience. We also noticed that the preference for the face-like stimulus was especially conspicuous in control females, which showed a remarkable preference level compared to all other groups (Supplementary Fig. S1; group mean preference index CTRL females 0.8029 [95% C.I. 0.6425 - 0.9632], one-sample t-test t_(13)_ = 4.081, uncorrected p = 0.0013; group mean preference index VPA females 0.4318 [95% C.I. 0.2064 - 0.6571]; t_(12)_ = 0.6419, uncorrected p = 0.5301; group mean preference index CTRL males 0.5711 [95% C.I. 0.3589-0.7832]; t_(18)_ = 0.7035, uncorrected p = 0.4907; group mean preference index VPA males 0.5163 [95% C.I. 0.3245 - 0.7081]; t_(18)_ = 0.1787, uncorrected p = 0.8602). However, given that no significant interaction between the factors emerged in our previous analysis, the gender effect observed requires extreme caution in the interpretation. Notably, regardless of the gender of the chicks examined, VPA-exposed chicks did not display any significant preference for the schematic face-like stimulus, indicating a detrimental effect of VPA on face-processing. Future studies will investigate the potential gender differences in the level of face-preference and in their susceptibility to VPA suggested by some of our data and clarify the mechanism of action of VPA on the development and expression of biological predispositions.

The increased latencies and the increased number of alternations observed in the VPA group, indicate that VPA exposure affects the visual preference for schematic face-configuration patterns without significantly hindering their capability for motoric activity during the test. In line with that, previous studies from our lab have shown that VPA exposure, at the dosage used in this study, does not significantly affect motor behaviour or discriminative abilities of simple artificial objects in domestic chicks (21).

A previous study has investigated the attentive behavior towards faces in VPA-exposed juvenile macaques (24). Using eye-tracking analysis to measure the animals’ attention, the authors found that juvenile VPA-treated monkeys spent significantly more time attending to nonsocial scenes than their control siblings. However, the study did not specifically investigate the predisposed response of visually naïve animals to faces compared to a visually equivalent stimulus without social content. In this respect, our study is the first to analyze a very early predisposed response for more strictly controlled stimuli in a visually naïve animal model of ASD.

## Conclusions

Altogether, this study and previous studies from our lab, demonstrate a detrimental effect of VPA, an anticonvulsant increasing the risk to develop ASD in humans, on the very early predisposed responses towards social stimuli in visually-naïve domestic chicks. Based on these results, we propose the domestic chicks as elective animal models to study these early-emerging neurobehavioural markers and to investigate the biological mechanisms underlying face processing deficits in ASD.

## Declarations

### Availability of data and materials

All data generated or analysed during this study are included in this published article (see supplementary information).

## Authors’ contributions

PS conceived and designed the experiments; AA and SP conducted the experiments; PS and OR-S analyzed the data; PS drafted the manuscript; AA, PS and OR-S wrote the manuscript. All authors read and approved the final manuscript.

## Acknowledgements

We thank Giorgio Vallortigara for his support, advice and comments on the manuscript. Dr. Tommaso Pecchia for help with the experimental apparatus, Grazia Gambardella for administrative help and Ciro Petrone for animal facility management.

## Figure Legends

**Figure S1. Social preference index of gender groups**. Bar graphs represent social preferences indexes for males and females. One-sample t-test on preference indexes indicate a significant difference from chance level for female control animals, but not for males or VPA-treated chicks of both genders. The number sign (^#^) indicate significant departures of the preference index from chance level (0.5), marked by the red line. Data represent Mean ± SEM, ^#^ uncorrected p < 0.05; ^##^ uncorrected p < 0.01.

